# The Cu-induced ScsD reductase is a membrane-bound redox partner of ScsB in the *Salmonella* envelope

**DOI:** 10.64898/2026.05.22.727170

**Authors:** Andrea A. E. Méndez, Juan J. Reinero, Zhenzhen Zhao, Bianca Bertonati, José M. Argüello, Fernando C. Soncini, Susana K. Checa

## Abstract

The intracellular environment *Salmonella* confronts during infection is characterized by multiple redox stressors including reactive oxygen species (ROS) and copper (Cu) ions. Under these conditions, alternative systems of thiol oxidoreductases such as the Cu induced Scs system are required to protect and repair periplasmic proteins. The *scsABCD* operon encodes three Dsb-like enzymes, ScsB, ScsC, and ScsD, and an accessory protein, ScsA. These proteins are required both for Cu resistance and H₂O₂ tolerance. ScsB and ScsC function analogously to the canonical DsbD/DsbC redox pair of thiol oxidoreductases. The absence of ScsC was shown to affect the folding/activity of periplasmic proteins involved in amino acid transport and redox homeostasis. Here, we focus in ScsD, the least characterized member of this system. Upon Cu-induced expression, ScsD localizes to the inner membrane, enabling its predicted C-terminal Dsb-like domain to be exposed to the periplasm. Functional analysis indicates that ScsD exists in a reduced state in the *Salmonella* envelope and serves as a redox partner of ScsB. ScsD exhibits *in vivo* disulfide reductase activity and restores a deficient disulfide reduction pathway in *Salmonella*. Similar to ScsC and ScsB, ScsD binds Cu(I) via the Cys residues of its Dsb-like domain; however, this metal interaction appears to lack relevance in Cu detoxification as no impact on intracellular Cu levels was observed. Our results define ScsD as a specialized membrane-bound thiol-disulfide reductase in the *Salmonella* envelope and highlight the versatility of the Scs system in maintaining periplasmic proteostasis when canonical pathways are compromised by host-imposed Cu stress.

**Importance:** Copper is a key component of the innate immune system, serving as a primary defense against pathogens like *Salmonella*. Copper overload targets the bacterial envelope, specifically attacking protein sulfhydryl groups. This causes protein misfolding and inactivation, disrupting essential processes like metabolism, transport and virulence. To survive this stress and restore thiol homeostasis, *Salmonella* utilizes the *scsABCD* operon. While the ScsB-ScsC redox pair is well-documented and some protein substrates identified, the role of ScsD remains undefined. This work characterizes ScsD as an inner-membrane-anchored thiol reductase and a new redox partner for ScsB. The ScsD/ScsB pair expands the bacterium’s protein quality control capacity, allowing *Salmonella* to maintain envelope homeostasis within the hostile, copper-rich environment of the host.

## INTRODUCTION

*Salmonella enterica* comprises a number food-borne, facultative intracellular pathogenic serovars that can cause diseases ranging from self-limiting gastroenteritis to severe invasive illness in susceptible hosts (1). During infection, this pathogen is exposed to multiple host-derived stressors. For instance, intestinal colonization triggers the production of reactive oxygen/nitrogen species (ROS/RNS) as part of the host defense response (2–4). Within phagocytic cells, in the lumen of endocytic vesicles, *Salmonella* encounters elevated concentration of copper ions (Cu), a toxic metal that induces redox stress and directly disrupts the structure and function of macromolecules, leading to further damage (5–8). These stressors primarily affect the bacterial cell envelope, a compartment essential for nutrients transport, metabolism, cell division and virulence (9–11). Accordingly, most components of the Cu resistance, redox defense and repair machineries are localized to the *Salmonella* envelope and play key roles in pathogenesis (12, 13).

Both ROS/RNS and Cu ions can alter the redox state of Cys residues of proteins. This modification frequently results in protein misfolding and/or inactivation (5, 10, 14, 15). Restoring redox homeostasis of the *Salmonella* envelope under these conditions, exceeds the capacity of the canonical periplasmic thiol oxidoreductases DsbA-DsbB and DsbC/G-DsbD. Under these conditions *Salmonella* deploys accessory systems such as ScsABCD. The Scs system is part of these protective networks, as its deletion compromises not only Cu and H_2_O_2_ tolerance, but also virulence (6, 16, 17). The *scsABCD* operon encodes a group of three Dsb-like enzymes (ScsB, ScsC and ScsD) and a putative accessory protein (ScsA) (18, 19). We previously demonstrated that expression of these proteins is induced by Cu stress and depends on CpxRA, a two-component regulatory system that coordinates much of the redox stress response at the bacterial envelope (6). ScsB is a large inner membrane (IM) protein that structurally resembles the canonical DsbD reductase. Both proteins differ mainly at their periplasmic thioredoxin-like α-domain, the region responsible for electron transfer to redox partners (20, 21). Notably, this domain is larger and structurally more complex in ScsB than in DsbD, suggesting the potential of ScsB for interaction with different partners.

Unidirectional electron transfer between the purified ScsB α-domain and the oxidized form of the periplasmic ScsC (22) suggested that these proteins form a redox pair analogous to the canonical DsbD-DsbC system. However, while the reductase/isomerase activity of DsbC is well established, the catalytic activity of ScsC is not. Both *in vivo* and *in vitro* studies suggested that ScsC may function as an oxidase or a reductase, depending on experimental conditions. Consistent with this, ScsC exhibits DsbD-dependent isomerase/reductase or oxidase activity when overexpressed in *Escherichia coli* Δ*dsbC* Δ*mdoG* or Δ*dsbA* mutant strains, an organism that lacks the *scs* locus (23). *In vitro*, both α-ScsB and ScsC can transfer Cu ions to CueP (22), a *Salmonella* periplasmic protein involved in Cu detoxification and required for virulence (13, 24). In terms of envelope homeostasis, deletion of *scsC* results in a reduced abundance of several periplasmic proteins involved in amino acid uptake and oxidative stress defense following Cu exposure (17). Surprisingly, CueP was not identified among these Cu-induced proteins.

In this work we focused on ScsD. Modeling using Alphafold-3 suggest that the 168 amino-acid polypeptide contains a thioredoxin-like domain (residues 36-163) anchored to the IM through a N-terminal hydrophobic α-helix (residues 1-31) (see Fig. 1B). The C-terminal region, likely exposed to the periplasm, harbors a putative CXXC catalytic motif as well as conserved residues present in IM-linked members of the TlpA thioredoxin-like superfamily, involved in cytochrome biogenesis (25–27) (Fig. S1). For instance, ScsD shared 36% amino acid identity with *Caulobacter crescentus* TlpA, and 30% with *Bradyrhizobium japonicum* TlpA and *E. coli* DsbE, and significant structure similarity (Fig. S1 and S2). We previously reported that Δ*scsC* and Δ*scsD* strains display similar sensitivity to Cu, whereas simultaneous deletion of both genes reduces Cu tolerance to levels comparable to those observed in a Δ*scsB* strain (6). In addition, both ScsC and ScsD provide significant protection against H_2_O_2_ stress (6). Based on these observations, and on the fact that TlpA and DsbE act as alternative substrates for ScsB and DsbD in *Caulobacter* or *E. coli,* respectively (20, 28), we hypothesize that ScsD functions as an alternative redox partner of ScsB.

**Figure 1.**
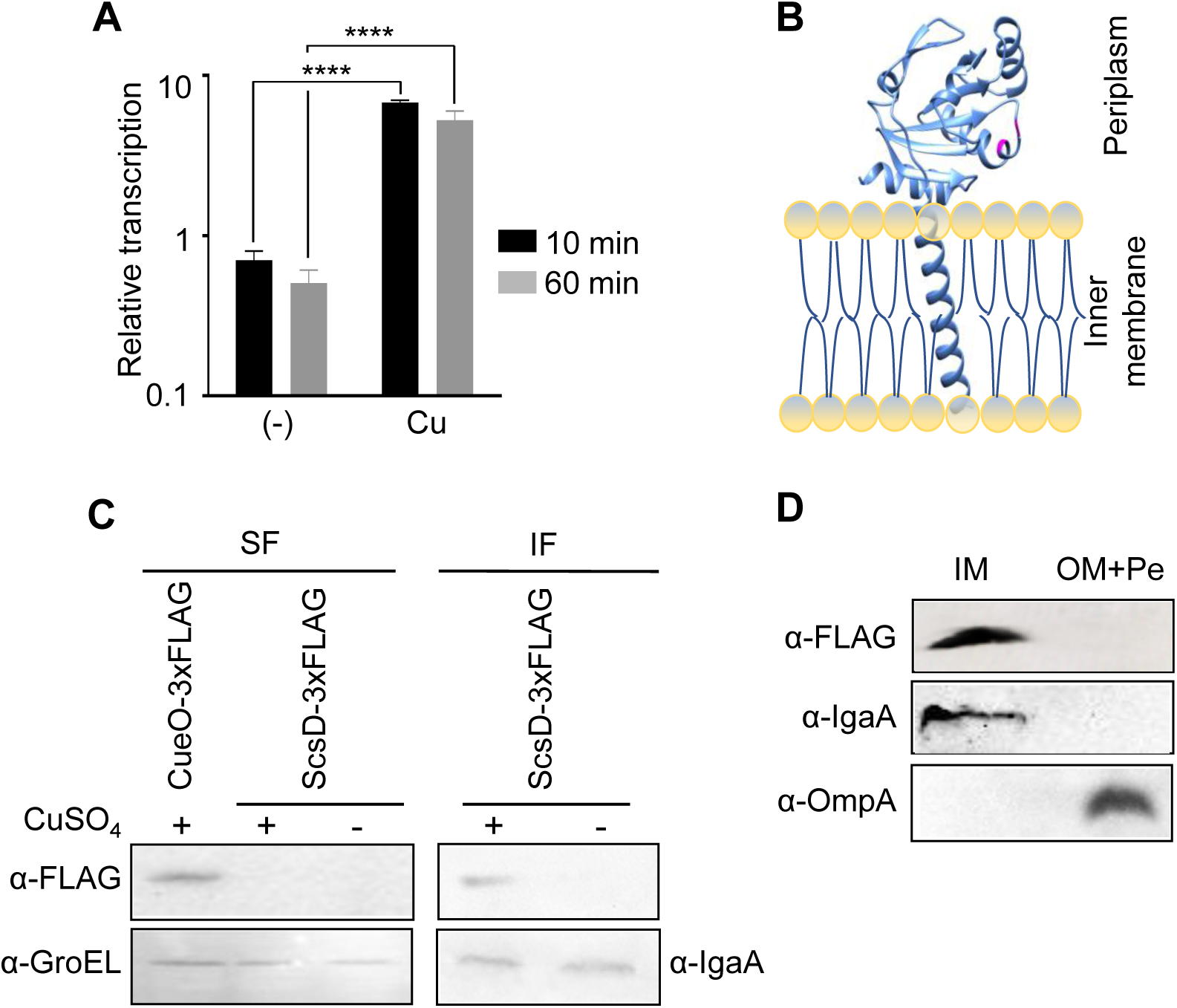
ScsD is a Cu-induced inner-membrane protein. (A) *ScsD* transcriptional profile upon Cu treatment. Relative transcription levels were obtained after normalizing the *scsD* values against those of *rnpB* and background (0 min) levels. Data are the mean ± SD, n=3. **** p <0.0001. (B) Schematic representation of the predicted ScsD intracellular location. Structural model of *S*. Typhimurium ScsD was obtained using AlphaFold 3. Cys71 and Cys74 are depicted in magenta. (C and D) SDS-PAGE 18% (w/V) and Western Blot analysis. In C, soluble (SF) or insoluble (IF) fractions were obtained from the ScsD-3xFLAG or CueO-3xFLAG 14028s strains grown overnight in the presence (+) or absence (-) of 1mM CuSO_4_. Immunodetection were done in parallel using primary antibodies directed against the Flag tag (α-FLAG) and GroEL (α-GroEL) or IgaA (α-IgaA) as loading controls of SF or IF, respectively. In D, the inner membrane (IM) fraction or in the combined outer membrane (OM) and periplasmic (Pe) fractions were obtained from the 14028s ScsD-3xFLAG strain grown in the presence of Cu ions. Immunodetection were done in parallel using α-FLAG and α-IgaA or the primary antibodies directed against OmpA (α-OmpA) as loading controls of the IM or the OM+Pe fractions, respectively.

Our results show that ScsD is a Cu-responsive, IM-anchored thiol-reductase that interacts with the α domain of ScsB via its C_71_GVC_74_ catalytic motif. Furthermore, ScsD can restore the loss of the canonical disulfide reduction pathway in the *Salmonella* envelope and likely cooperates with ScsB to preserve homeostasis under Cu-induced redox stress.

## RESULTS

### ScsD is an inner-membrane protein expressed in response to Cu

ScsD contributes to Cu tolerance in *S. enterica* (6). Cu-dependent induction of the *scsD* gene was initially inferred from transcriptional profiling of the 5’ regions upstream of the first two genes in the *scs* operon, *scsA* and *scsB* (6). To validate this observation, we quantified *scsD* transcript levels following Cu exposure. *scsD* expression increased rapidly after Cu addition and remained unchanged after 60 min (Fig. 1A).

Cu-dependent induction and intracellular localization of ScsD were analyzed by western blot using a chromosomal *scsD*-3xFLAG tagged strain. Notice that addition of the 3xFLAG tag in the chromosomal copy of *scsD* did not affect ScsD folding or activity, since similar levels of Cu tolerance were observed in either the *scsD-*3xFLAG containing or the WT strains (Fig. S3A). In accordance, expression of the tagged protein from a multicopy plasmid complemented the Cu sensitive phenotype of the Δ*scsD* strain as well as the native protein (Fig S3B).

The chromosomally encoded ScsD-3xFLAG protein was detected in the presence of Cu and exclusively in the insoluble fraction (Fig. 1C). Further cell fractioning indicated its association with the cell inner membrane (Fig. 1D). A strain expressing the soluble (periplasmic) Cu responsive CueO-3xFLAG protein (13) was included as control. The use of antibodies against the cytoplasmic GroEL chaperon, the IM-associated protein IgaA or the outer-membrane linked OmpA porin supported our observations (Fig. 1D) and confirmed its proposed location based on *in silico* predictions of ScsD structure (Fig. 1B).

### ScsD exists in a reduced state in the *Salmonella* envelope

ScsD has only two cysteine residues (C_71_ and C_74_), both part of a putative catalytic motif (Fig. 1B and S1). To evaluate the reactivity of these residues, we cloned, expressed and purified the soluble periplasmic domain of ScsD (ScsDp). Far-UV circular dichroism of purified proteins (Fig. S4B and S4D) showed secondary structures (∼35% α-helix, ∼30% β-barrel and ∼5% random coil) in agreement with the Alphafold-3 structural model (Fig 1B). Purified ScsDp was incubated with and without DTT and then subjected to chemical modification with 4-acetamido-4‘-maleimidostilbene-2,2‘-disulfonate (AMS). Reacting with free thiols, a AMS molecule increases the molecular mass of the protein by 490 dalton (29). SDS-PAGE analysis revealed a protein shift corresponding to the ScsDp-AMS adducts only in samples pre-incubated with DTT (Fig. 2A). This suggested that sulfhydryl groups of the purified ScsDp are oxidized and only upon DTT treatment they became reduced to accept the alkyl group.

**Figure 2.**
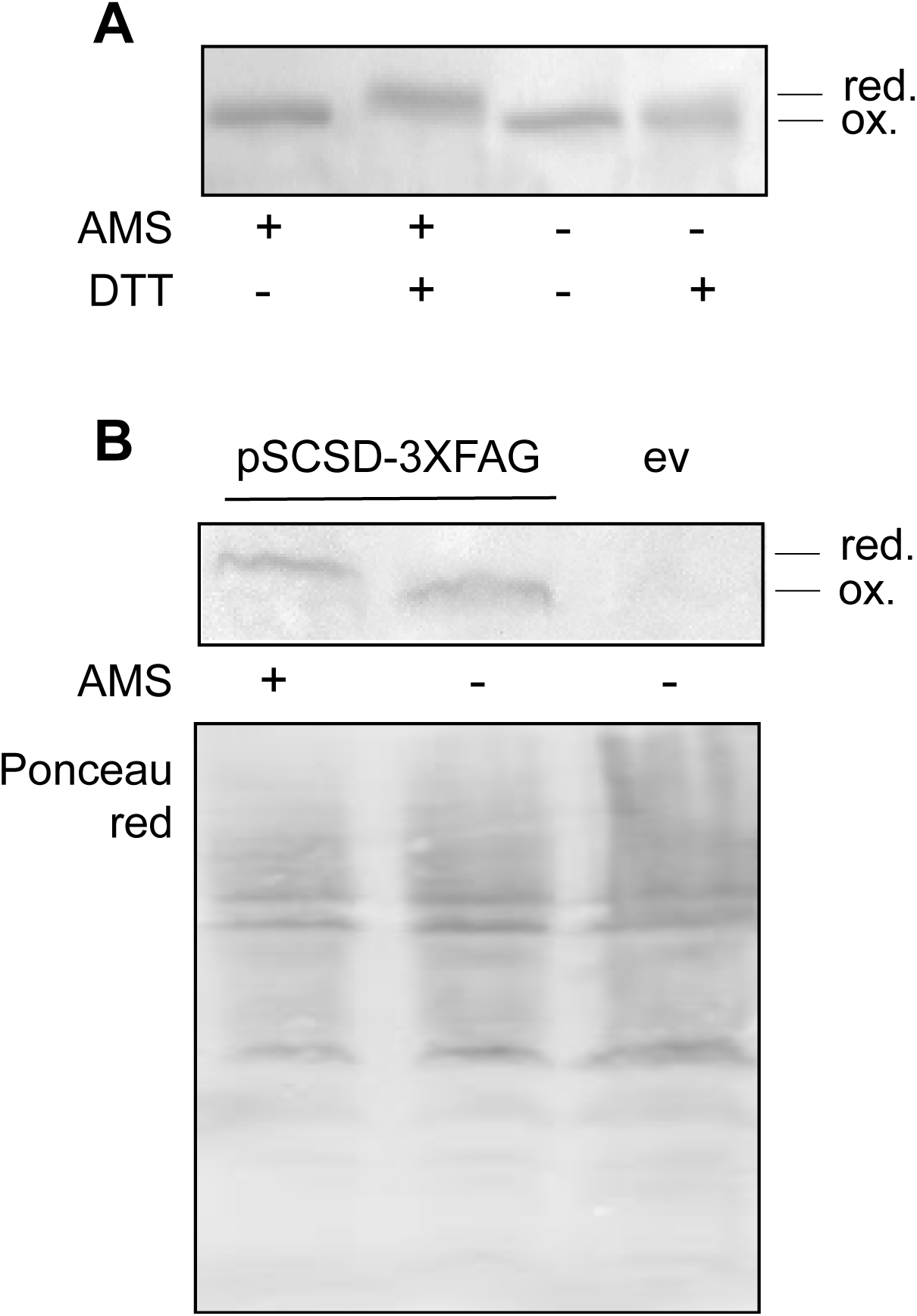
Redox state of ScsD. (A) Non-reducing SDS-PAGE (18% w/V) of purified ScsDp treated with AMS and/or DTT, as indicated. (B) Whole cell lysates obtained from 14028s carrying pSCSD-3xFLAG or the empty vector (ev) were incubated for 2 h to induce protein expression and treated or not with AMS, as indicated. Non-reducing SDS-PAGE and Western Blot were done using α-FLAG monoclonal antibody. Ponceau red staining is shown as loading control. The reduced (red.) or oxidized (ox.) state of the analyzed protein is indicated.

The *in vivo* redox state of ScsD was determined using a similar AMS modification assay applied to cell extracts. A strain expressing the fully functional ScsD-3xFLAG from a plasmid (Fig. S3B) was grown to exponential phase, followed by protein precipitation with trichloroacetic acid and subsequent AMS treatment. Expression of the tagged protein was performed from a heterologous promoter, rather than from its Cu-inducible chromosomal promoter, minimizing potential interference arising from Cu-dependent redox activity. The formation of ScsD-3xFLAG-AMS adducts indicates that ScsD is predominantly in a reduced state in the *Salmonella* envelope (Fig. 2B), with the sulfhydryl groups of the C_71_GVC_74_ motif available to interact with putative redox partners or with Cu ions.

### ScsD is a ScsB redox partner

The structural and biochemical similarities of ScsD and ScsC, together with their additive participation in *Salmonella* tolerance to Cu and H_2_O_2_ (6), led us to investigate whether ScsD is a ScsB redox partner. We used the redox-trap strategy, a method previously employed to test the participation of different Dsb-like proteins (21), to evaluate the formation of ScsDp/ScsBα like complexes. The Cys_74_Ser ScsDp and the Cys_113_Ala 6xHis-ScsBα mutant forms of the proteins were obtained and purified (Fig. S4 and S5A). These protein variants, named here m-ScsDp and m-ScsBα for simplicity, have the second and the first Cys in their active sites replaced by Ser or Ala, respectively (Fig. 3A). If ScsD and ScsB function as redox partners, the presence of these mutations would allow the formation of a S-S bridge between the remaining Cys while blocking the electron transfer needed to resolve and release the individual proteins. The complexes formed by combining these purified mutant proteins can be detected by non-reducing SDS-PAGE. As controls, we included the purified wild-type forms of both ScsDp and ScsBα, as well as ScsC, a bona fide periplasmic redox partner of ScsBα (17) and its mutant Cys_69_Ser version (m-ScsC) (Fig. S3 and S4).

**Figure 3.**
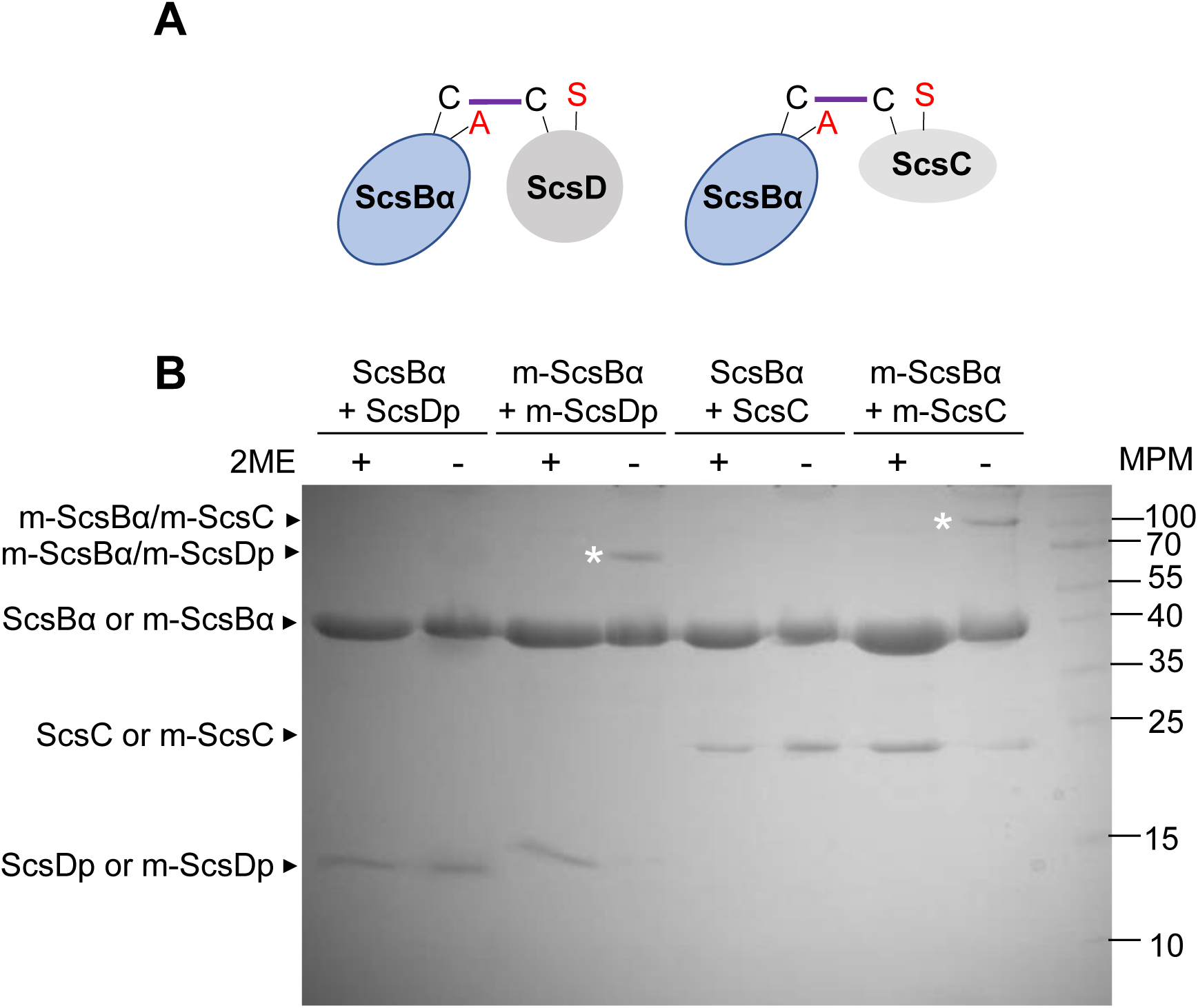
ScsD and ScsC are ScsB redox partners. (A) Schematic representation of the redox trap strategy. The mutated residue in each protein is shown in red and the covalent interaction between them represented by a curve line. (B) Non-reducing SDS-PAGE (18% w/V) analysis of trapped redox protein complexes. The purified proteins included in each mix as well as addition/not addition of β-mercaptoethanol (2ME) are detailed. The bands corresponding to the individual proteins, the covalent complexes (*) or the protein marker employed are indicated. The m-prefix refers to Cys_113_Ala-ScsBα, Cys_74_Ser ScsDp Cys_69_Ser-ScsC.

A higher molecular weight band corresponding to the trapped redox complexes was observed in samples containing m-ScsBα and either m-ScsDp or m-ScsC (Fig. 3). Addition of β-mercaptoethanol to these samples resulted in disruption of the S-S bond and release of the individual proteins composing each redox pair (Fig. 3B). As expected, the bands corresponding to the redox complexes were not observed when the wild-type form of the partner proteins were mixed, both in the presence or absence of the reductant (Fig. 3B). The redox-trap strategy applied here not only allowed us to propose ScsD as a new ScsB partner, but also to corroborate the ScsC-ScsBα interaction, which was previously predicted based on electron transfer studies (22).

### ScsD harbors disulfide reductase activity and can restore the loss of disulfide reduction pathway in *Salmonella*

*S. enterica* serovar Typhimurium (*S.* Typhimurium) 14028s strain forms large, wrinkled colonies with the characteristic *rdar* (red, dry, and rough) morphotype (Fig. 4A) when grown on salt-free LB (SLB) agar plates supplemented with Coomassie Brilliant Blue and Congo Red (CR plates) (30, 31). Deletion of the canonical periplasmic disulfide oxidase DsbA results in smaller, thicker colonies with intensely colored center compared to the wild type (30) (Fig. 4A). To assess the *in vivo* catalytic activity of ScsD, we first investigated its ability to suppress the Δ*dsbA* morphotype. Expression of ScsD from the pSCSD plasmid failed to restore the wild type *rdar* morphotype, yielding colonies that remained phenotypically similar to the Δ*dsbA* strain (Fig. 4A). This contrasted with the phenotype exhibited by the complemented strain. Notice that SscD expressed from pSCSD was confirmed to be functional as it fully complemented the Cu sensitivity phenotype of a Δ*scsD* strain (Fig. S5). These results suggest that ScsD lack disulfide oxidase activity *in vivo*.

**Figure 4.**
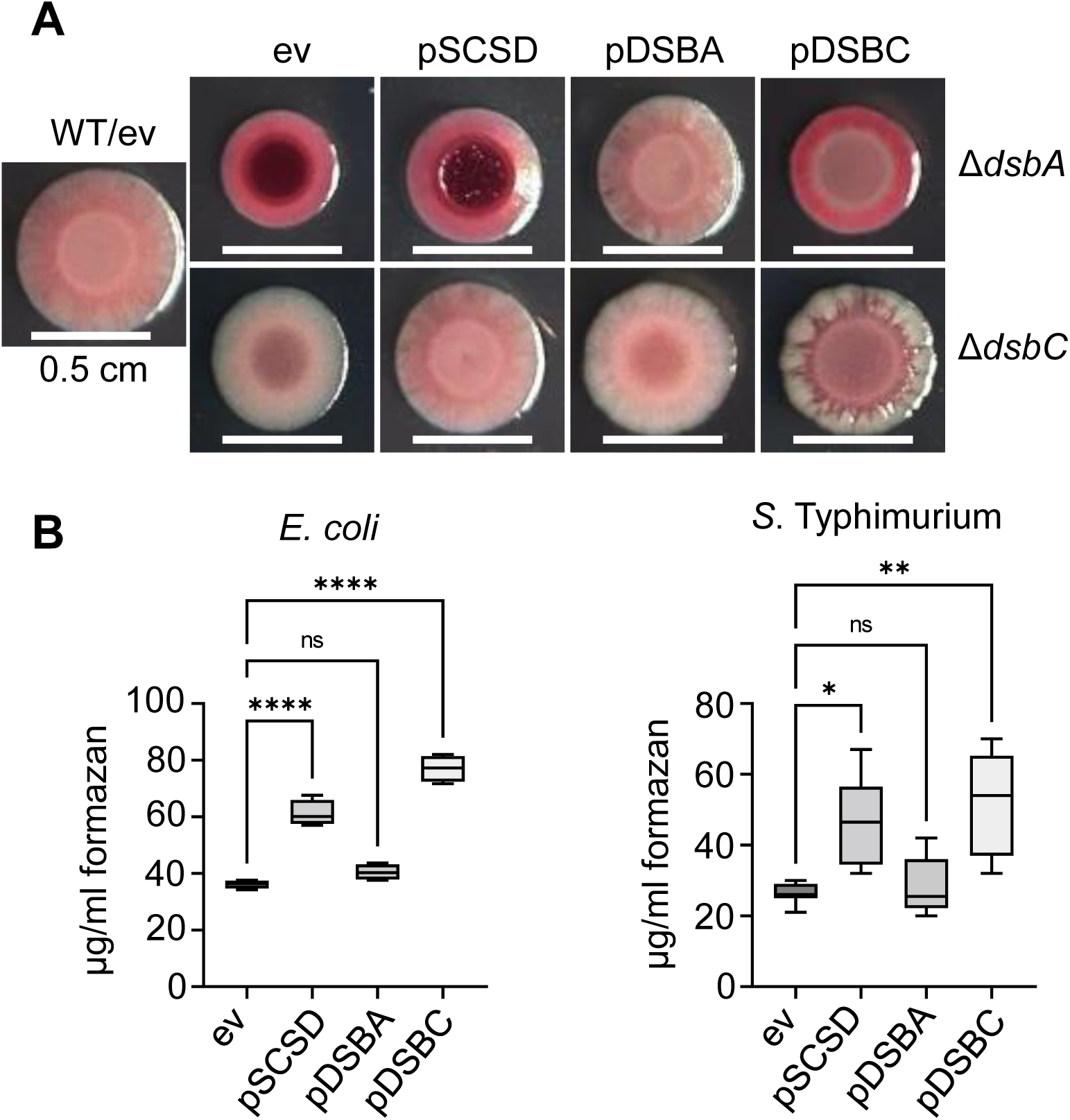
ScsD functions as a periplasmic reductase. (A) *Rdar* colony phenotype of the wild-type 14028s (WT) strain or the Δ*dsbA* or Δ*dsbC* mutants carrying expression plasmids for the *Salmonella* ScsD (pSCSD), DsbA (pDSBA), DsbC (pDSBC) or the empty vector (ev) (see Table S1). All strains were grown on CR agar plates at 28 °C for 72 hours. (Scale bars, 0.5 cm.). (B) Nitro Blue Tetrazolium reduction assay done in *E. coli* BL21(DE3) strain or *S.* Typhimurium 14028s carrying the indicated plasmids. Formazan production was evidenced spectrophotometrically at 572 nm after 45 min incubation with NBT in a 37 °C water bath. Data are the mean ± SD, n=4.0. ns, not significant; *, p < 0.05; **, p <0.01; ****, p <0.0001.

Interestingly, the Δ*dsbA* strain overexpressing the reductase/isomerase DsbC also failed to complement *rdar* morphotype, although these colonies exhibited a different color distribution compared to those carrying the empty vector or pSCSD (Fig. 4A). This led us to evaluate the *rdar* morphotype of the Δ*dsbC* (Fig. 4A), since no prior influence of DsbC on biofilm formation has been reported. After 72 h of growth at 28 °C, the Δ*dsbC* mutant produced colonies similar in size but with a smoother and paler appearance compared to the wild type. Overexpression of ScsD in the Δ*dsbC* background restored the wild type morphotype. In contrast, the strain expressing DsbA from the plasmid failed to restore the wild-type appearance, while the Δ*dsbC*/pDSBC strain exacerbated the *rdar* phenotype (Fig. 4A). Overall, these results showed that ScsD can functionally substitute the DsbC reductase/isomerase activity under these conditions.

Further evidence for the catalytic activity of ScsD was obtained by exploiting its ability to reduce nitroblue tetrazolium (NBT) to formazan that can be monitored by absorbance at 572 nm (32). Overexpression of ScsD in *E. coli* resulted in an almost two-fold increase in NBT reduction, levels like those obtained in the strain carrying pDSBC (Fig. 4B). In contrast, the strain overexpressing DsbA exhibited no detectable activity above the empty-vector control. When the assay was performed in the *S.* Typhimurium strain, a similar trend was observed (Fig. 4B). These results supported the reductase activity proposed for ScsD.

### ScsD coordinates Cu(I)

Cu binding capacity was previously reported for both ScsC and ScsBα, and proposed, although not demonstrated, to be linked to their CX_2/3_C motif (22). Here, ScsDp/Cu binding stoichiometry was analyzed by AAS. The isolated soluble domain was incubated under saturating conditions of Cu (10:1, Cu:protein) and the free metal removed by chromatography. Under these conditions, ScsDp bounds ∼1 equivalent of Cu (Fig. 5A).

**Figure 5.**
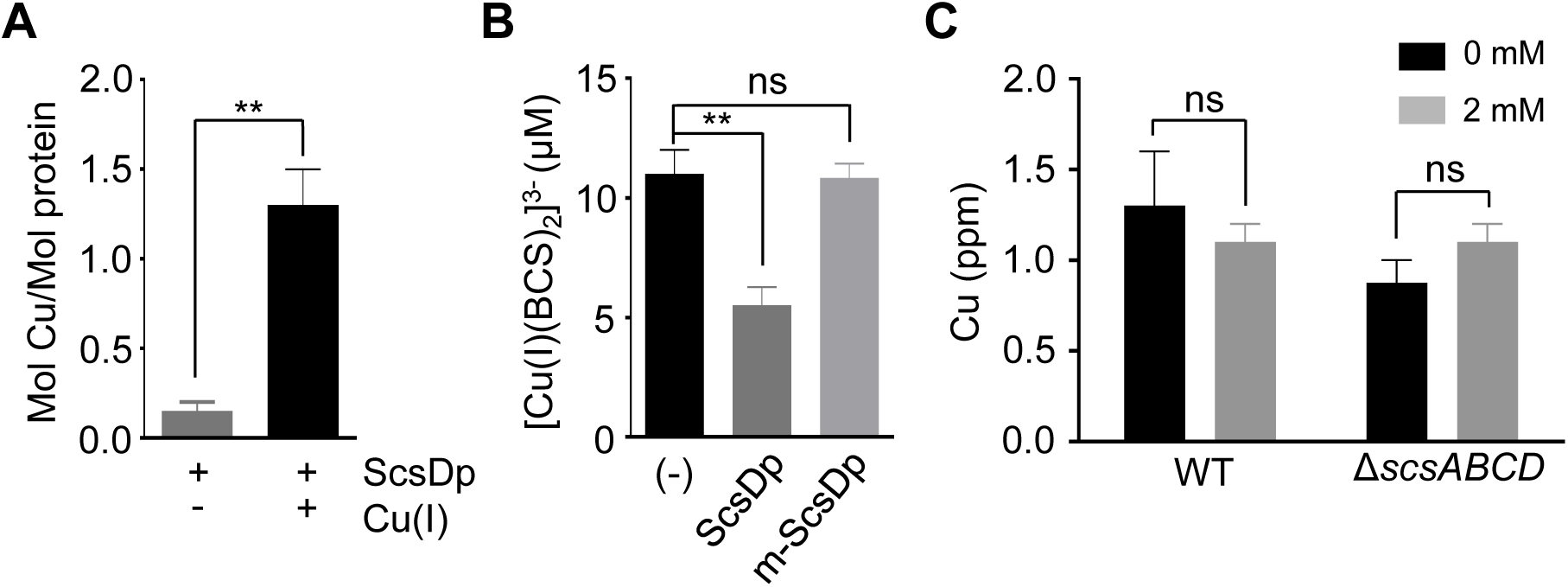
ScsD binds Cu(I) *in vitro* but this does not affect intracellular Cu levels. (A) Purified protein ScsDp (10 nM) was incubated with/without CuSO_4_ for 10 min at room temperature. Bound Cu(I) was determined by AAS and normalized against the amount of protein in the sample. (B) ScsDp or m-ScsDp (10 µM) was incubated with 25 μM BCS and 10 μM CuSO_4_ during 5 min at room temperature. [Cu(I)(BCS)_2_]^3-^ complex was quantified photometrically. (C) Copper uptake from 14028s WT and Δ*scsABCD* strains after 60 min incubation in 0 o 2 mM CuSO_4_. Data are the mean ± SD, n=3.0. ns, not significant; **, p <0.01.

To further evaluate the interaction of ScsDp with Cu, we performed competition binding assays using the specific Cu(I) chelator bathocuproinedisulfonic acid disodium salt (BCS). When present in equimolar amounts to the [Cu(BCS)_2_]^3-^ complex, purified ScsDp was able to displace approximately 50% of the bound Cu(I) (Fig. 5B). This indicates that, under these conditions, ScsDp competes effectively with BCS for the metal ion. Given that the stability constant for the Cu(BCS)_2_ complex is β= 10^19.8^ M^−2^, the estimated apparent dissociation constant (*K_d_*) for the ScsDp-Cu(I) complex is in the femtomolar (10^-15^ M) range. This high affinity for Cu-binding is similar to that reported for ScsC (22). On the other hand, no competition was observed in samples containing the chelator and the same concentration of m-ScsDp (Fig. 5B), suggesting that at least the Cys74 residue of ScsD is essential for Cu(I) coordination.

Although ScsD, ScsC and ScsB are Cu-binding proteins, their role in Cu homeostasis seems not to be associated with Cu chelation or transport, since deletion of the whole *scsABCD* operon had little or no effect on the intracellular Cu levels (Fig. 5C). Altogether, these results indicate that both the ScsC/ScsB and ScsD/ScsB complexes are rather involved in preserving the redox homeostasis in conditions of Cu stress than in metal handling.

## DISCUSSION

*Salmonella* encodes a diverse repertoire of periplasmic Dsb-like oxidoreductases that are essential for bacterial fitness and virulence (16, 19). Like other bacterial pathogens, its interaction with the host is strongly influenced by transition metals, particularly Cu (33). When present in excess, Cu ions readily coordinate with thiol groups, leading to protein misfolding or displacement of other metals from their native binding sites, ultimately resulting in protein inactivation (33). In addition, Cu can trigger or exacerbate oxidative stress, thereby amplifying cellular damage. The *Salmonella scsABCD* operon, which is absent in *E. coli* but conserved among other enteropathogens, encodes three Dsb-like proteins that collectively protect against Cu and redox stress and contributes to virulence (19).

Characterization of the Scs system has focused primarily on the IM-bound ScsB and the periplasmic protein ScsC, which has been proposed as its redox partner (20, 21). We previously showed that third Dsb-like component, ScsD, contributes additively with ScsC to Cu tolerance, as the Δ*scsC* Δ*scsD* double mutant is as sensitive to Cu as the Δ*scsB* strain (6). In this study, we demonstrate a redox interaction between ScsD and ScsB and provide evidence for ScsD subcellular localization, expression, and function in the *Salmonella* periplasm.

Although *scsD* is the last gene in the operon, its transcription is strongly induced by Cu (Fig. 1) (6). This induction parallels the behavior of other periplasmic copper-responsive genes, such as *cueO* and *cueP*, reinforcing the view that ScsD and its partners are part of the *Salmonella* copper-stress stimulon (33). As ScsC and ScsB, ScsD interacts with Cu using its catalytic C_71_GVC_74_ motif (Fig. 5A). However, deletion of the entire *scs* operon did not alter intracellular Cu levels, even under Cu stress conditions (Fig. 5C and data not shown). These observations and the *in vivo* disulfide reductase activity of ScsD (Fig. 4), indicate that this redox enzyme, like the other Scs proteins, is not directly involved in Cu handling, but rather in the protection and repair of thiol groups in periplasmic proteins during Cu stress.

Cell fractionation and Western blot analysis of chromosomally encoded ScsD (Fig. 1C and D), together with the production and purification of its functional C-terminal soluble domain (ScsDp) (Fig. S4A and B), corroborate the *in silico* predictions. ScsD is anchored to the inner membrane via its N-terminal helix, while exposing its globular thioredoxin-like domain containing the catalytic C_71_GVC_74_ motif to the periplasm (Fig. 1B). *In vivo* AMS-shift assays further indicated that the Cys residues of ScsD are maintained reduced within the *Salmonella* envelope (Fig 2B). Consistent with this, reductase activity assays (Fig. 4B) demonstrated that ScsD exhibits catalytic properties comparable to those of DsbC, the canonical enterobacterial disulfide reductase/isomerase (34). Functional restoration of biofilm formation by ScsD in a Δ*dsbC* background, but not in the Δ*dsbA* strain (Fig 4A), reinforces both its reductase activity and its physiological relevance as a component of the periplasmic redox homeostasis machinery.

Given that DsbC requires DsbD for its reductase activity but not for its isomerase function (19), it is reasonable to infer that ScsD likewise depends on an associated electron donor *in vivo*. By using redox-trap experiments (Fig. 3), we demonstrated that ScsD forms a covalent complex with ScsB, providing direct biochemical evidence for the existence of a functional redox pair between these enzymes. The ability of ScsB to deliver electrons to either ScsC and ScsD would allow to broad the spectrum of substrates processed by this system, a feature shared by several Dsb-like redox networks (35).

As mention in the introduction, ScsD structurally resembles TlpA from *B. japonicum* (Fig. S2), an IM-anchored, periplasmic-facing thiol/disulfide oxidoreductase that, like ScsD, participates in Cu homeostasis (27). In this bacterium, TlpA is essential for maintaining the reduced state of the membrane-bound periplasmic metallochaperone Scol (36). This reduction enables ScoI to bind Cu and subsequently transfer it to the active site of heme-Cu cytochrome oxidase subunit II (CoxB), which itself also requires TlpA-mediated reduction to accept the metal. Although *Salmonella* lacks ScoI or CoxB homologs, it encodes the periplasmic Cu chaperone CueP that, like the Scs system, is absent in *E. coli* and its expression is controlled by the CpxRA system (12). Similar to ScoI, CueP requires reduced Cys residues to coordinate Cu ions (37, 38). While ScsC has been proposed as the CueP specific reductase, several observations argue against this model: CueP was not identified among ScsC redox substrates (17), no direct redox interaction between CueP and ScsC has been detected *in vitro*, and ScsC efficiently transfers Cu(I) to the reduced form of CueP rather than reducing it (22). Together, these findings suggest that ScsD may represent the missing CueP reductase required for metalation of this Cu-chaperone.

The opportunistic pathogen *C. crescentus* encodes a functional ScsB homolog that forms a redox pair with ScsC (20, 39), but lacks ScsA or ScsD (19). In this organism, a TlpA homolog, structurally similar to ScsD (Fig. S2), also forms a redox pair with ScsB (20). The substrate of the alternative ScsB/TlpA pathway is PprX, a periplasmic peroxiredoxin involved in redox homeostasis. Considering that *Salmonella* ScsA is a putative lipoprotein required for H_2_O_2_ detoxification (16, 19), contains six Cys residues, and has homology to peroxiredoxins, we hypothesize that this protein may be an additional ScsD substrate. Other potential ScsD substrates may include membrane-associated proteins whose thiol groups are damaged during Cu stress.

Our results demonstrate that ScsD is a copper-induced, IM-anchored reductase that functions as another ScsB redox partner. We propose that ScsD/ScsB and ScsC/ScsB cooperate to prevent the accumulation of misoxidized or metal-poisoned envelope proteins during Cu stress, thereby contributing to protein quality control and promoting *Salmonella* survival within the host environment.

## MATERIALS AND METHODS

### Bacterial strains and growth conditions

Bacterial strains used in this study are listed in Table S1. Cells were grown overnight at 37°C in Luria-Bertani Broth (LB) or minimal medium M9, supplemented with 25 mg ml^-1^ kanamycin (Km),10 mg ml^-1^ chloramphenicol (Cm), 100 mg ml^-1^ ampicillin (Amp) or 100 mg ml^-1^ streptomycin (Sm) when required. CuSO_4_ salt used were of ACS analytical grade ≥98.0% purity.

### Genetic and molecular biology methods

Oligonucleotides used in these studies were provided by Macrogen and are listed in Table S2.

Chromosomal gene insertion of the 3xFLAG tag was carried out as previously described (13). Briefly, a 1530 pb fragment carrying 3xFLAG-*Km*^R^ cassette was amplified using Q5 High-Fidelity DNA polymerase (New England Biolabs) and oligonucleotides ScsD-3xFLAG Fw and ScsD-3xFLAG Rv (Table S2) from the pSUB11 plasmid (Table S1). The resulting amplicon was introduced into the *S.* Typhimurium 14028s chromosome by one-step linear transformation followed by Lambda Red-mediated recombination (40). Km resistant colonies were selected and proper insertion verified by colony-PCR.

pSCSD complementation plasmid (Table S1) was constructed using the T5 exonuclease–dependent assembly (TEDA cloning) protocol (41). The *scsD* gene was amplified from the chromosome of *S.* Typhimurium 14028s strain and inserted into the pUHE21-2lacI*^q^* plasmid (pSCSD) (41). For generation of pScsD-3xFLAG, the *scsD-3xFLAG* fragment was amplified from PB15186 strain (Table S1), digested with *BamH*I and *Sal*I and cloned into the pUC19 plasmid digested with these enzymes.

To generate the expression plasmids for the soluble periplasmic domain of ScsD (ScsDp, residues 36-163), ScsC without its signal peptide (residues 25-186) or the ScsBα domain (residues 30-270) the Restriction Free (RF) method (42) were applied. These gene fragments were amplified from the chromosome of *S.* Typhimurium 14028s strain (Table S1) using the Q5® High-Fidelity DNA polymerase and specific oligonucleotides listed in Table S2. The purified products corresponding to *scsDp* or *scsC*, or to *scsBα* were used as mega-primers in RF reaction with the pQE80L-*6xHis*-*mbp*-*gfp* or pET32a-*6xHis*-*gfp* vectors (Table S1), respectively (43). The remaining non-amplified vector was eliminated by adding 20U of *Dpn*I. Digestion was stopped by heat-inactivation at 65°C for 10 min. The mutant version of ScsDp (Cys_74_Ser-ScsDp or m-ScsDp), ScsC (Cys_69_Ser-ScsC or m-ScsC) and ScsBα (Cys_133_Ala-ScsBα or m-ScsBα) were generated by applying site directed mutagenesis to the plasmids carrying the wild-type copies. Complementary primers carrying the desired point mutation were designed (Table S2) and used in a Q5-mediated amplification followed by digestion with *Dpn*I to eliminate the wild-type copy of the plasmid.

All the plasmids were initially introduced into competent *E. coli* TOP10 cells. Positive Ampicillin resistant clones were verified by colony PCR and DNA sequencing.

### RNA isolation and analysis of gene expression by qRT-PCR

*Salmonella* WT LB cultures at an optical density at 600 nm (OD_600_) of 0.5 were exposed to 2 mM CuSO_4_ for 10 or 60 min. RNA from three biological replicates was pooled to constitute a single RNA sample for sequencing. RNA extraction was done as previously described (31). Total RNA samples were treated with RQ1 RNase-free DNase I (Promega). RNA integrity was verified by agarose electrophoresis. cDNA was synthesized with MoMLV-reverse transcriptase (Promega). Quantitative real-time PCR was done in an Accurate 96 Real-Time PCR System (DLAB), using the HOT FIREPol® EvaGreen® qPCR Mix Plus (ROX) (Solis BioDyne) and the primers listed in Table S2. Cycling parameters were: denaturation at 95 °C for 12 min; 40 cycles of 95 °C for 10 s, 60°C for 20 s and 72 °C for 20 s; and a final extension of 72 °C for 10 min. Expression was calculated by 2^−ΔΔCT^ method (31), using *rnpB* as a housekeeping gene. Each cDNA sample was run in technical triplicate and repeated in at least three independent sets of samples.

### Cell fractionation and Immunoblot analysis

*Salmonella* cell cultures at OD_600_ of 0.5 were treated with 1 mM CuSO_4_ for 18 h, harvested by centrifugation and re-suspended in 1 mL of Tris-HCl (pH 8), 10 mM EDTA, 1 mM phenylmethylsulfonyl fluoride (PMSF) buffer. After sonic disruption (30% amplitude) on ice for 2 min, with on/off intervals of 2 s (44), the homogenates were centrifuged (12,000 g, 30 min, 4 °C), soluble fraction collected, and pellet resuspended in Tris-HCl 10 mM (pH 8) EDTA 1 mM buffer for analysis.

To obtain the fractions corresponding to the IM and the OM plus periplasm, collected cells were resuspended in a lysis buffer containing sucrose 20% (w/v), 0.5 mM EDTA and 0.15 mg ml^-1^ lisozyme and incubated on ice for 40 min. MgCl_2_ at 20 mM was added and samples were centrifuged 20 min 11.200 g to separate a supernatant including the periplasm (Pe) and the outer membrane (OM) fraction from the pellet containing intact protoplasts. Further sonic disruption and centrifugation were done to discard the cytoplasm and recover the inner-membrane (IM) in the pellet (45).

Protein concentration was measured using Pierce™ BCA Protein Assay Kit. All fractions were separated in 18% (w/v) SDS-PAGE and electro-transferred to nitrocellulose membranes and treated with the indicated primary antibodies followed by either rabbit or mouse secondary antibody conjugated with horseradish peroxidase (HRP) (13). Mouse anti-FLAG (α-FLAG) monoclonal antibodies were for sigma Sigma-Aldrich. Rabbit polyclonal antibodies directed against GroEL, IgaA and OmpA were from Laboratory stocks. Immunoreactive bands were visualized using the SuperSignal® West Femto Maximum Sensitivity Western Blotting Substrate (Thermo Scientific™).

### Expression and purification of recombinant proteins

*E. coli* clones expressing the fusion proteins 6xHis-MBP-ScsDp, 6xHis-MBP-ScsC or their mutant versions were grown in LB broth supplemented with 100 μg ml^-1^ of ampicillin. At the optical density at 600 nm (OD_600_) of 0.5, 0.75 mM isopropylthio-β-galactoside (IPTG) was added to induce expression. Incubation was allowed to proceed for additional 5 hours at 26 °C. Cells were harvested by centrifugation and re-suspended in 10ml of a lysis buffer containing 50 mM Tris-HCl (pH 8,0), 0.3 M NaCl, 10 mM imidazole, 2 mM EDTA, 10% glycerol, and 1 mM PMSF. Cell suspensions were sonicated and centrifuged as described above to collect the soluble fractions. After filtration using a 0.2 µm filter, these fractions were loaded into amilose resin columns (New England Biolabs) for purification according to manufacturerś instructions. As the fusion proteins contains the TEV recognition sequence followed the 6xHis-MBP region, we treated the purified fusion proteins with the protease in a 1:100 TEV:protein ratio for 3 hr at room temperature in a buffer containing 50 mM Tris-HCl (pH 8.0), 0.5 mM EDTA and 1 mM dithiothreitol (DTT). HisPur Ni-NTA resin columns (Thermo Scientific) were used to separated purified ScsDp from 6xHis-MBP and the 6xHis-TEV, following manufactureŕs instructions.

The 6xHis-ScsBα protein or its mutant version m-ScsBα were purified from *E. coli* extracts prepared as described above and purified using HisPur Ni-NTA resin columns.

Purity of the resulting proteins was verified by 12% (w/v) SDS-PAGE (Fig. S3 and S4). ScsDp and m-ScsDp structural integrity was also analyzed by circular dichroism (Fig. S3) in a Jasco J-810 spectropolarimeter (JASCO, Easton, MD) with a 1 mm path length cuvette. The experiments were carried out at 25°C for the near UV (250-320nm) region. Spectrum was measured in triplicate at a scan speed of 20 nm min^-1^, with a time constant of 1 s. Data were averaged to minimize noise, and molar ellipticity calculated as described (44) using the Dicroprot® software.

### Sulfhydryl reactivity

For *in vitro* sulfhydryl reactivity, purified ScsDp was reduced or not with 1 mM DTT in 20 mM Tris-HCl (pH 8.0), for 1 h on ice. AMS (4-Acetamido-4′-maleimidostilbene-2,2′-disulfonate; Sigma Aldrich) was prepared in 50 mM Na_3_PO_4_ buffer (pH 7.0), and added to a final concentration of 10 mM. The reaction was carried out for 10 min at room temperature. Protein modification was performed in a buffer containing 50 mM Tris-HCl (pH 7.5), 50 mM NaCl and 1 mM EDTA, using protein samples at a concentration of 0.5 mg ml^-1^. The reaction was stopped by lowering the pH with 0.1 volume of 3 M HCl, after which the proteins were analyzed by nonreducing SDS-PAGE on 18% (v/W).

For *in vivo* pScsD-3xFLAG analysis, fresh overnight cultures were diluted 1/20 in 10 ml of M9 minimal media supplemented with Amp in 100 mL flasks and incubated at 37 °C with shaking at 220 rpm. After 1 h, the OD_600_ was measured and adjusted to 1 unit using fresh media, prior to precipitation with 10% trichloroacetic acid (TCA) and incubation on ice for 1 h (46). The precipitates were centrifuged for 15 min at 16,000 *g* at 4 °C, washed with cold acetone and resuspended in 100 µL of 10 mM AMS in 50 mM Tris-HCl (pH 8.0) 1% SDS and 1 mM EDTA. The mixes were vortexed 30 min at room temperature followed by 5 min incubation at 37 °C, centrifuged, and the supernatants discarded. Finally, the pellets were solubilized in 100 µl of Laemmli sample buffer without reducing agents, separated in a nonreducing 18% (w/v) SDS-PAGE and analyzed by Western blot using α-FLAG antibodies.

### *In vitro* redox trap assay

Complex formation between ScsBα and either ScsC or ScsDp was evidenced through trapping mixed disulfide intermediates between the mutant forms of these proteins, as previously described (21). Briefly, m-ScsDp or m-ScsC were mixed with 6XHis-mScsBα at 1:3 ratio in 50 mM Tris-HCl (pH 8.0) 0.3 M NaCl, 250 mM imidazole and 10% glycerol. Samples containing the wild-type form of each pair were processed in parallel to serve as controls. All protein mixtures were incubated at 37°C for 24 h. Aliquots (100 µl) were mixed with SDS-PAGE loading buffer containing or not β-mercaptoethanol and resolved by 12%(w/v) SDS-PAGE prior to photographical recording.

### Determination of *in vivo* ScsD redox activity

The catalytic activity of ScsD was assessed based on its ability to complement the deficient biofilm formation of Δ*dsbA* or Δ*dsbC* mutant *Salmonella* strains (30). The *rdar* morphotype was judged visually on CR plates (31). One µl of bacterial cultured grown at OD_600_ of 5.0 in LB were spotted onto the surface of agar plates containing LB without salt (SLB) supplemented with 40 mg l^-1^ Congo red, 20 mg l^-1^ Coomassie brilliant blue G-250, 100 mg ml^-1^ Amp and 0.75 mM of IPTG. Plates were incubated at 28°C for 72 hours and development of the colony morphology and color analyzed.

Nitroblue tetrazolium (NBT) reduction assays (32) were also used to test ScsD activity *in vivo*. A single colony of each strain was grown overnight in 1 ml of LB broth supplemented with 100 µg ml^-1^ Amp and 1 mM IPTG to induce recombinant proteins expression. Cultures were adjusted to an OD_600_ of 0.5 and incubated with 0.05 % NBT in Hank‘s solution (BDH Chemicalb Ltd, Poole, England) for 45 min at 37 °C. The reaction was stopped with 0.25 M HCl, samples centrifuged at 1000 g for 15 min. Supernatants were discarded and pellets were resuspended with 0.5 ml dimethylsulphoxide (DMSO). Formazan was measured spectrophotometrically at 572 nm.

### Copper tolerance

Functionality of ScsD or its 3XFLAG tagged form expressing from its chromosomal copy or from a plasmid were assessed by testing Cu tolerance on the wild-type or the Δ*scsD Salmonella* strains as previously described (6, 13). Briefly, a 10 µl drop of a 7-fold dilution of each bacterial culture was applied in the surface of LB agar plates supplemented with the indicated concentrations of CuSO_4_. Plates were incubated for 24 h at 37°C under aerobic condition after photographic recording.

### Copper binding assay

To determine the total amount of Cu bound to ScsDp, 10 nM of the purified protein was incubated with a 10-fold molar excess of CuSO_4_ in 25 mM HEPES (pH 8.0), 150 mM NaCl and 10 mM ascorbate for 10 min at room temperature with gentle shaking. The unbound Cu was removed using a Sephadex G-10 column (Sigma). After one step washing, the protein was concentrated using a 3-kDa Centricon. The amount of Cu bound to ScsDp was determined by atomic absorption spectroscopy (AAS), as described (47).

The interaction of ScsDp with Cu ions was assessed by competition assays using the chromogenic Cu(I)-binding compound bathocuproinedisulfonic acid (BCS) (47). This and the use of the mutant form of the protein allow us to estimate affinity and the residues involved in the interaction. Reduced apoproteins were obtained by incubation with 5 mM DTT for 1 h on ice. Excess DTT was removed by buffer exchange into freshly prepared 25 mM HEPES (pH 8), 150 mM NaCl and 10 mM ascorbic acid using Centricon Amicon® Ultra-4 (Millipore). ScsDp or m-ScsDp (10 µM) was incubated with 10 μM CuSO_4_ and 25 μM BCS in a buffer containing 25 mM HEPES pH 8, 150 mM NaCl and 10 mM ascorbic acid for 5 min at room temperature with gentle agitation. Then, absorbance of the [Cu(I)(BCS)_2_]^3-^ complex was measured at 483 nm (BCS ɛ = 13,000 M^−1^ cm^−1^) (47).

### Whole-cell Cu uptake assays

Fresh overnight cultures of the WT and Δ*scsABCD* strains were diluted 1:100 into 10 mL of LB medium supplemented with Amp (100 μg ml^-1^) in flasks and incubated at 37 °C with shaking at 220 rpm. When the cultures reached an OD₆₀₀ of 0.5, CuSO₄ was added to a final concentration of 2 mM, and incubation was continued for 60 min. Cells were harvested by centrifugation (10,000 g for 2 min at 4°C) and pellets were washed twice with ice-cold buffer (50 mM Tris-HCl pH 7,5, 150 mM NaCl). Cells were mineralized using fuming HNO_3_ (trace metal grade) at 80°C for 60 min and, after cooling, treated with 2 M H_2_O_2_ for complete digestion. If necessary, samples were diluted with ddH_2_O before AAS determination of Cu levels.

### Bioinformatics and data processing

3D structure prediction of proteins was carried out using AlphaFold3 Server (alphafoldserver.com). All structures were visualized and modeled in ChimeraX-1.7.1 (19). Sequence alignment of ScsD orthologs was done with Clustal Omega (48). Deconvolution of circular dichroism data was done using the Dicroprot® software.

Data visualization and statistical analyses were performed using GraphPad Prism version 9.0 (GraphPad Software, San Diego, CA). Differences between groups were analyzed using a one-way ANOVA followed by Tukey’s multiple comparisons test. A p-value <0.05 was considered statistically significant.

## ACKNOWLEDGMENTS

We thank Jimena Zoni for technical Assistance. This work was supported by CONICET (PIP-600 to S.K.C), Agencia I+D+I (PICT 2021-0779 to A.A.E.M.) and NIH (R01A1150784 to J.M.A.). These funders had no role in study design, data collection and interpretation, or the decision to submit the work for publication. A.A.E.M. is a CONICET postdoctoral fellow. S.K.C and F.C.S. are career investigators of CONICET. F.C.S. is also a Career Investigator of Consejo de Investigaciones de la Universidad Nacional de Rosario.

## AUTHOR CONTRIBUTIONS

Andrea A.E. Mendez: Data curation, Conceptualization, Formal analysis, Investigation, Methodology, Validation, Visualization, Funding acquisition, Writing – original draft, Writing – review & editing. Juan J. Reinero: Data curation, Formal analysis, Investigation, Methodology, Validation, Visualization, Writing – original draft. Zhenzhen Zhao: Data curation, Formal analysis, Investigation, Methodology, Validation, Visualization. Bianca Bertonati: Data curation, Formal analysis, Methodology, Validation, Visualization. José M. Argüello: Funding acquisition, Project administration, Resources, Formal analysis, Investigation, Supervision, Visualization, Writing – review & editing. Fernando C. Soncini: Conceptualization, Data curation, Formal analysis, Project administration, Resources, Supervision, Writing – original draft, Writing – review & editing. Susana K. Checa: Funding acquisition, Project administration, Resources, Investigation, Validation, Visualization, Formal analysis, Supervision, Writing – original draft, Writing – review & editing.

## Supplementary Information

**Figure S1.**
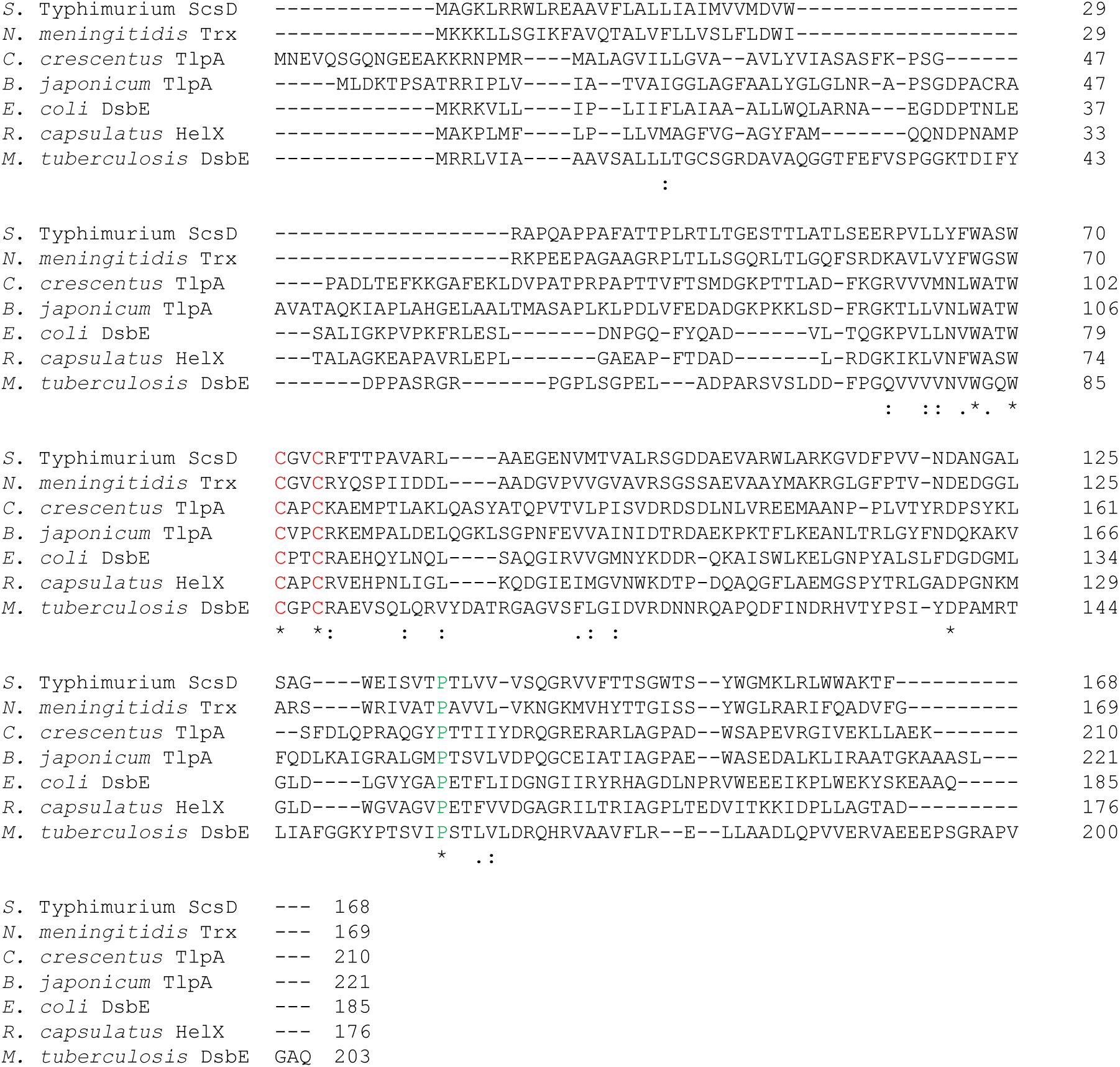
Protein sequence alignment of *Salmonella* ScsD with homologues of the TlpA thioredoxin-like superfamily using Clustal Omega. Marked in red are Cys residues forming the CXXC catalytic motif and in green a highly conserved Pro residue, reported to be essential for protein structure in *E coli* DsbE (26). Analyzed proteins and their percentage of identity with ScsD from *S.* Typhimurium (A0A0F6AZT7) are: a putative thioredoxin (Trx) from *Neisseria meningitidis* (Q9JXX7), 46%; TlpA from *Caulobacter crescentu*s (A0A290MK97), 36%; TlpA from *Bradyrhizobium japonicum* (WP_014491649.1), 31%; DsbE from *Escherichia coli* (P0AA86), 30%; HelX from *Rhodobacter capsulatus* (P36893), 29%; DsbE from *Mycobacterium tuberculosis* (A0A0E9AGM2), 28%. The code between parenthesis is the uniprot.org accession number.

**Figure S2.**
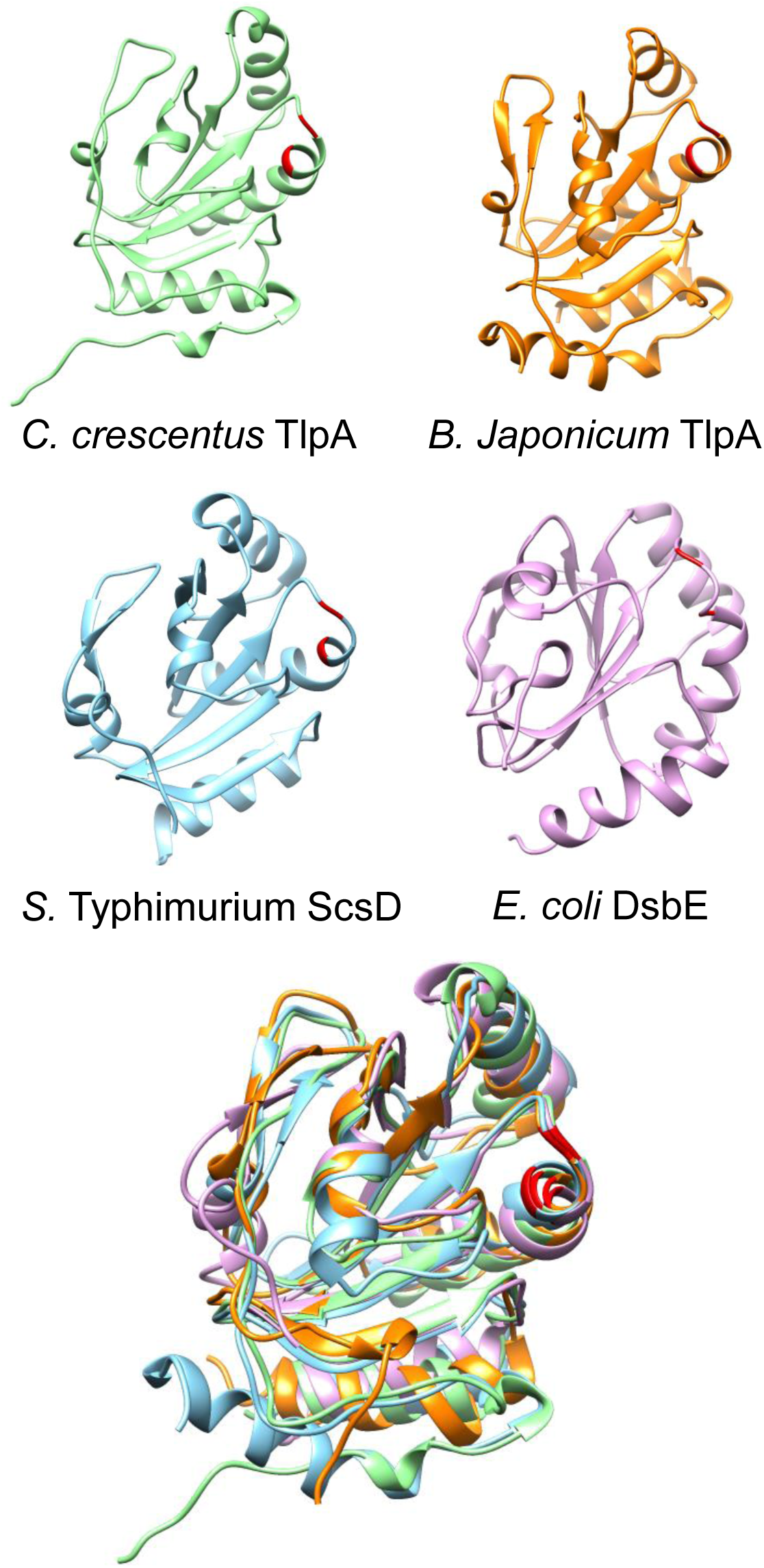
ScsDp and its structural homologues. The 3D structures of *C. crescentu*s TlpA (AlphaFold3), *B. japonicum* TlpA (PDB: 1JFU), *E. coli* DsbE (PDB: 3K8N) and *S.* Typhimurium ScsD (AlphaFold3) were analyzed using the ChimeraX-1.7.1 software. Multiple structure alignment is also shown. The catalytic Cys residues are highlighted in red.

**Figure S3.**
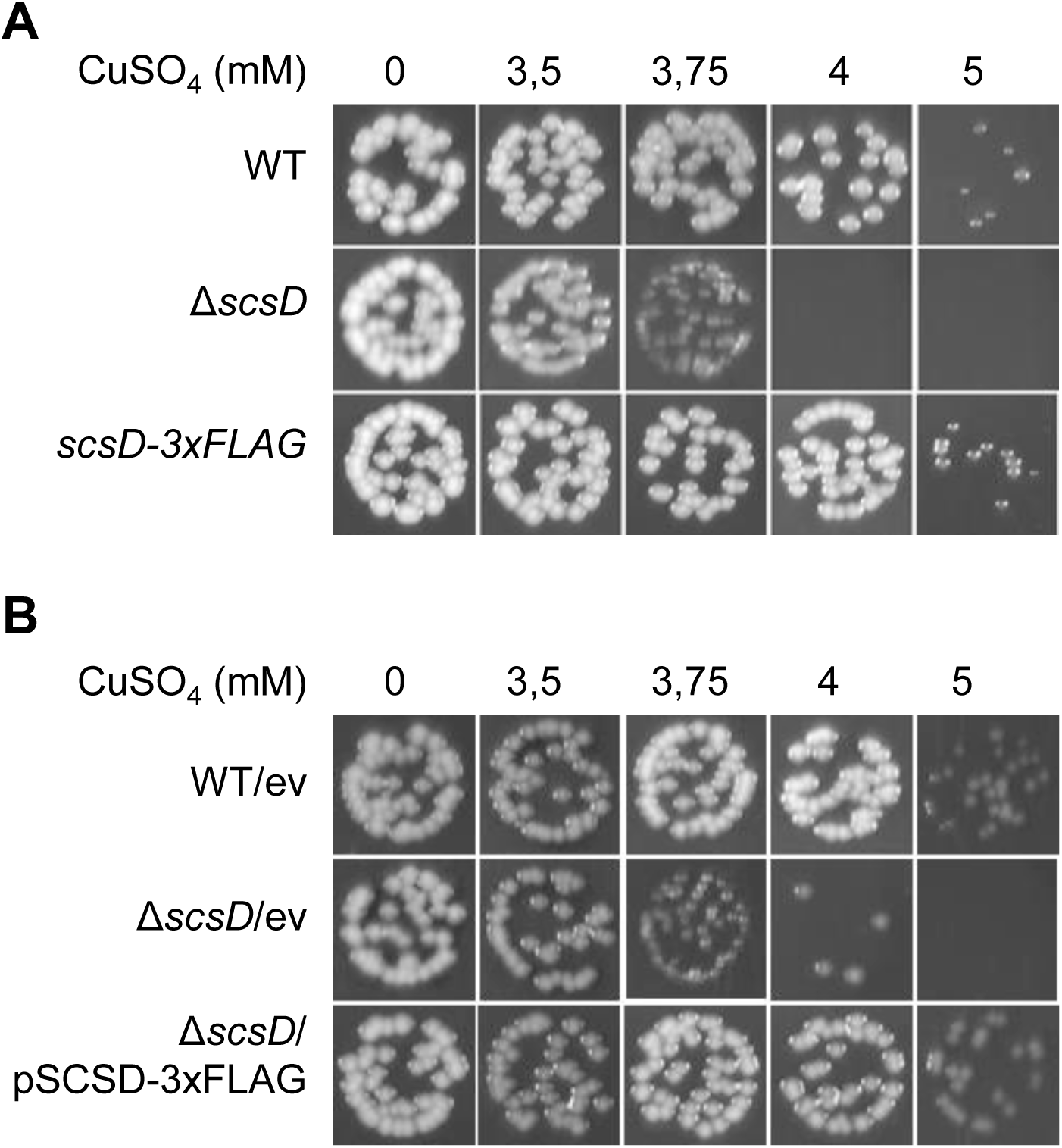
Both chromosomal and plasmid-derived ScsD-3xFLAG can restore Δ*scsD* mutant sensitivity to Cu. The Cu tolerance profiles of wild-type (WT), Δ*scsD* and chromosomal ScsD-3xFLAG *S.* Typhimurium strains (A) or the WT or the Δ*scsD* carrying the empty vector (ev) or the pSCSD-3xFLAG plasmid (B) were compared. All strains were grown in Luria-Bertani broth plates containing increasing amounts of CuSO_4_ in aerobic conditions. After incubation at 37°C for 24h, the plates were photographed. The data are representative images of at least three independent experiments done in duplicate.

**Figure S4:**
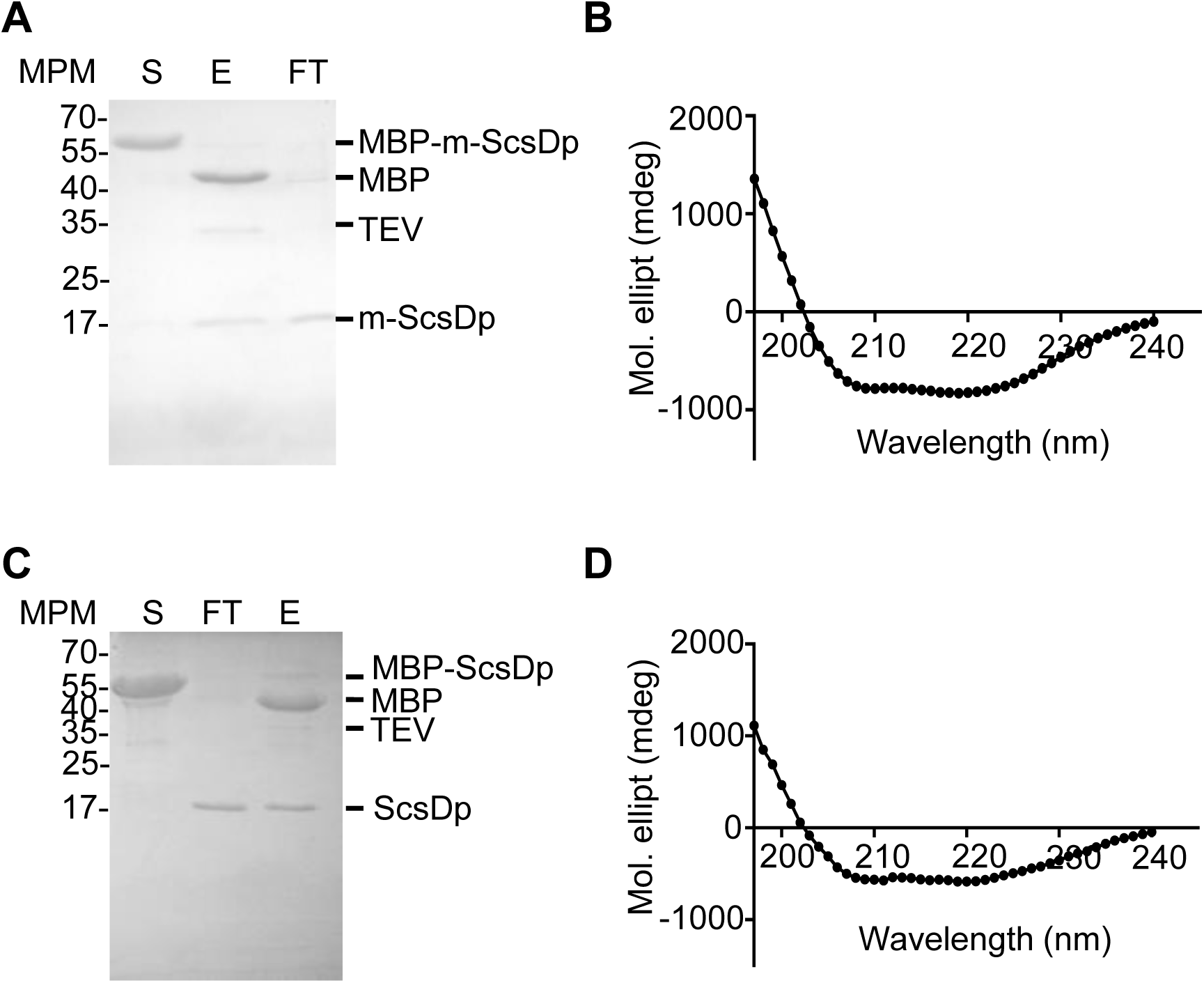
Purification and analysis of m-ScsDp and ScsDp. (A and C) Coomassie Blue-stained SDS-PAGE 18% (w/v) of purified 6xHis-MBP-m-ScsDp or 6xHis-MBP-ScsDp before or after cleavage by 6xHis-TEV protease and subsequent purification with Ni-NTA resin. S is the purified fusion protein (sample) applied to the column; E, elution using 500 mM imidazole buffer; FT, flow through using a 20 mM imidazole containing buffer. The proteins corresponding to each band are indicated. (B and D) Far UV circular dichroism of purified m-ScsDp or ScsDp, respectively.

**Figure S5.**
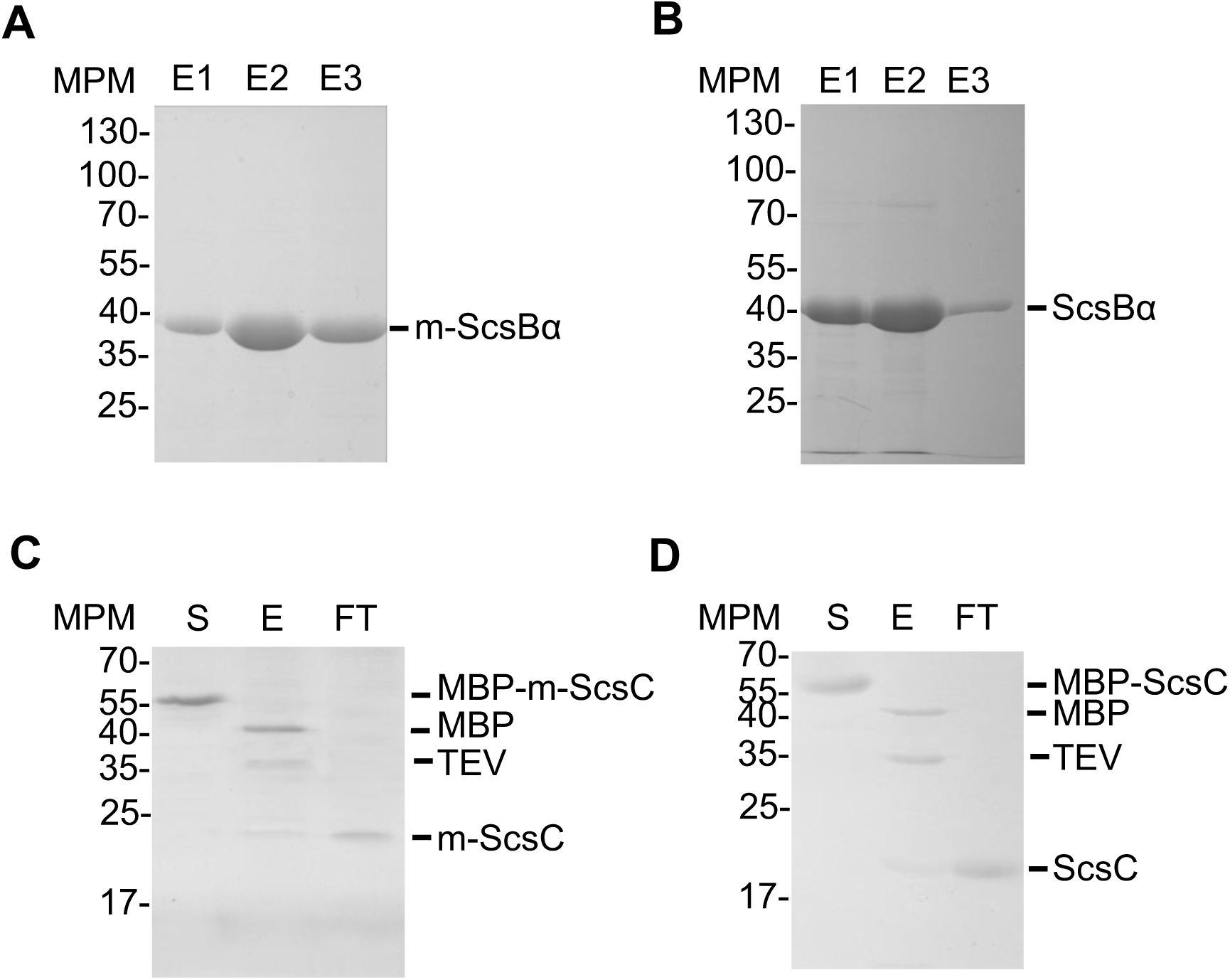
Purification of 6xHis-ScsBα, ScsC and their mutant forms. Representative Coomassie Blue-stained SDS-PAGE 12% (w/v) of the 6xHis-m-ScsBα (A) or 6xHis-ScsBα (B) purified using Ni-NTA columns. E1-3: fractions eluted with 250 mM imidazole buffer. (C and D) Coomassie Blue-stained SDS-PAGE 18% (w/v) of purified 6xHis-MBP-m-ScsC or 6xHis-MBP-ScsC before or after cleavage by 6xHis-TEV protease and subsequent purification with Ni-NTA resin. S is the purified fusion protein applied to the column; E, elution using 500 mM imidazole buffer; FT, flow through using a 20 mM imidazole containing buffer. The proteins corresponding to each band are indicated.

**Figure S6.**
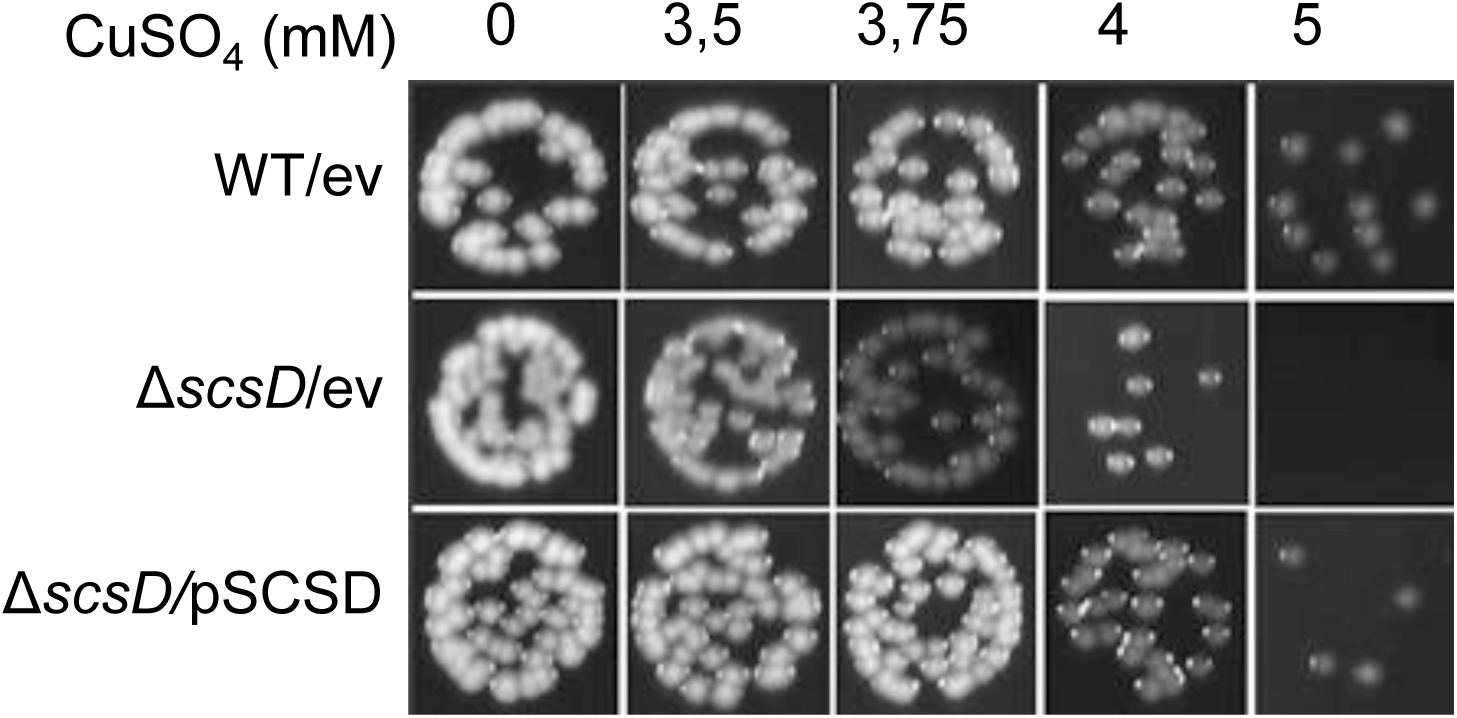
ScsD expressed in trans complements the Δ*scsD* Cu sensitivity. Comparative Cu tolerance profiles of the WT strain or the Δ*scsD* mutant carrying the empty vector (ev) or the ScsD expression plasmid (pSCSD). Bacterial growth was analyzed on LB broth plates containing increasing amounts of CuSO_4_. After incubation at 37°C for 24h under aerobic conditions, the plates were photographed. The data are a representative image of at least three independent experiments done in duplicate.

---

**Table S1.**
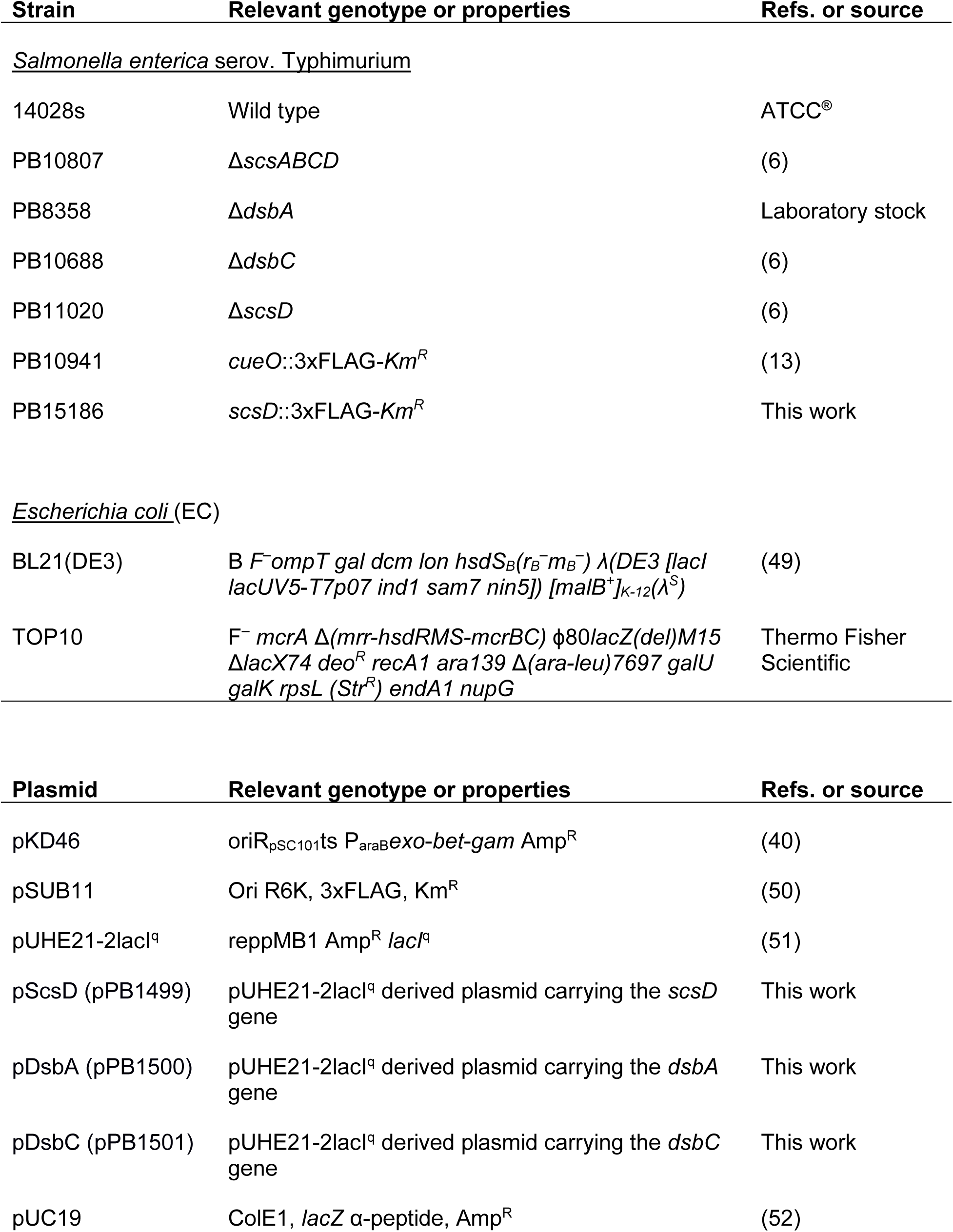
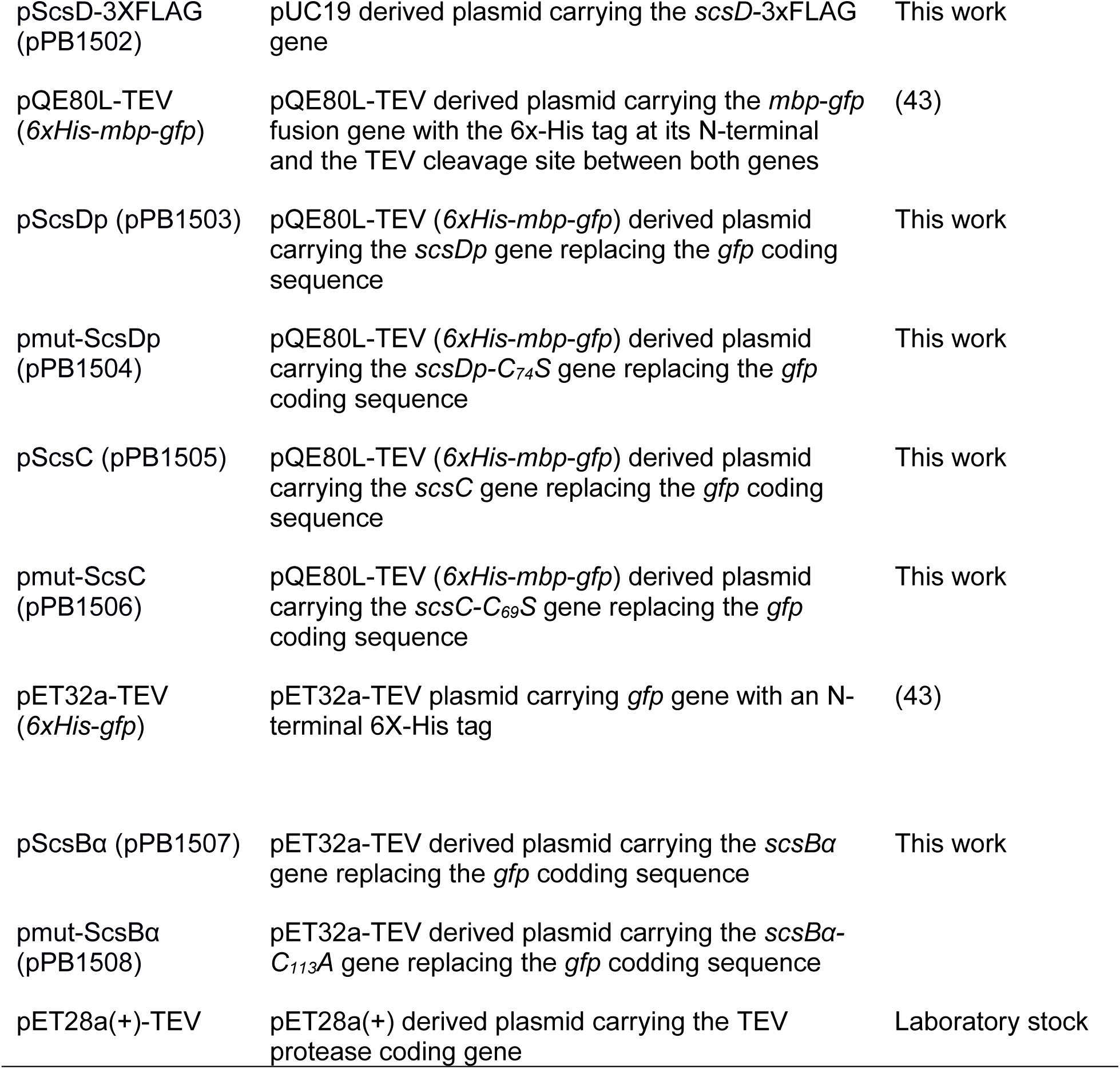
Bacterial strains and plasmids used in this study.

**Table S2.**
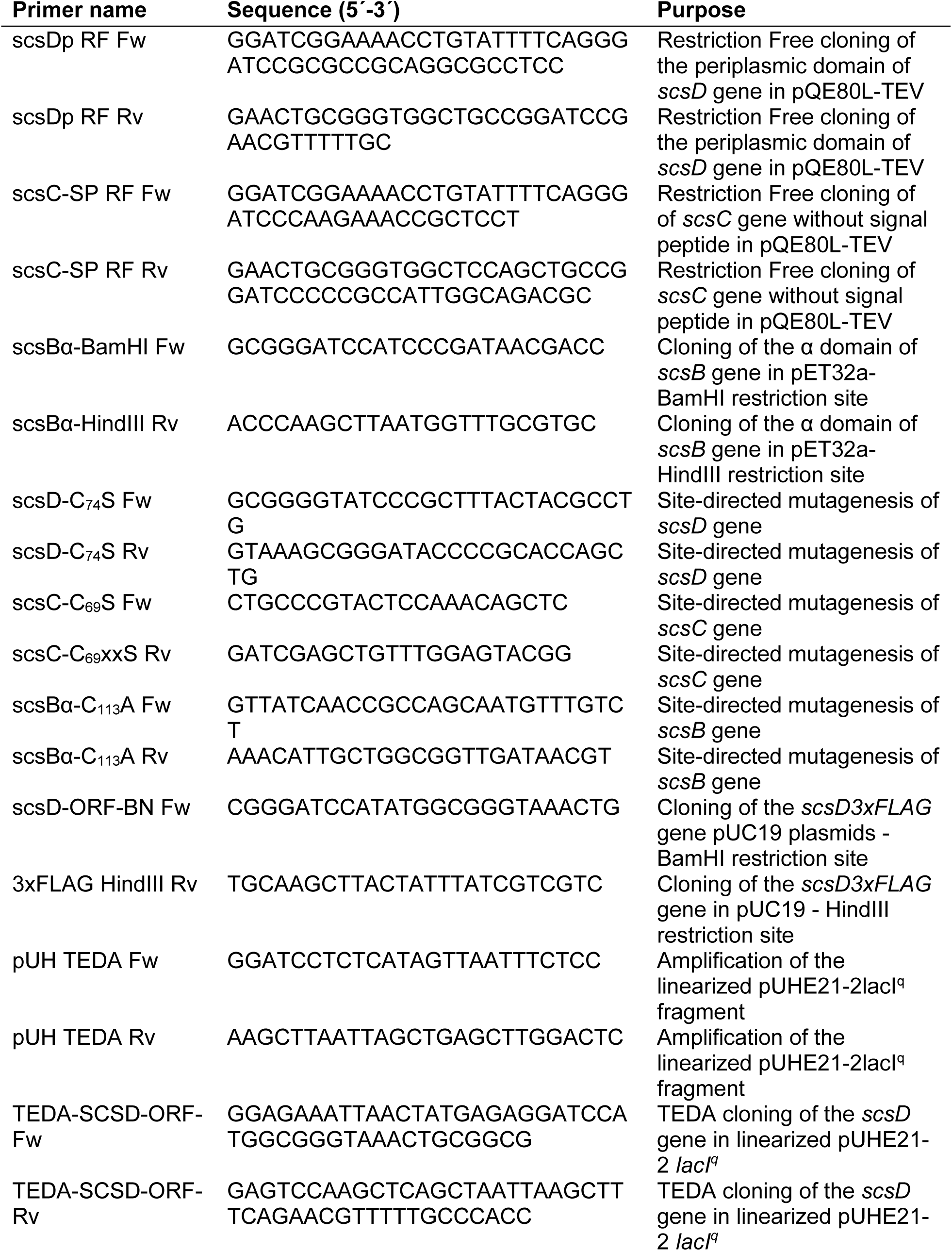
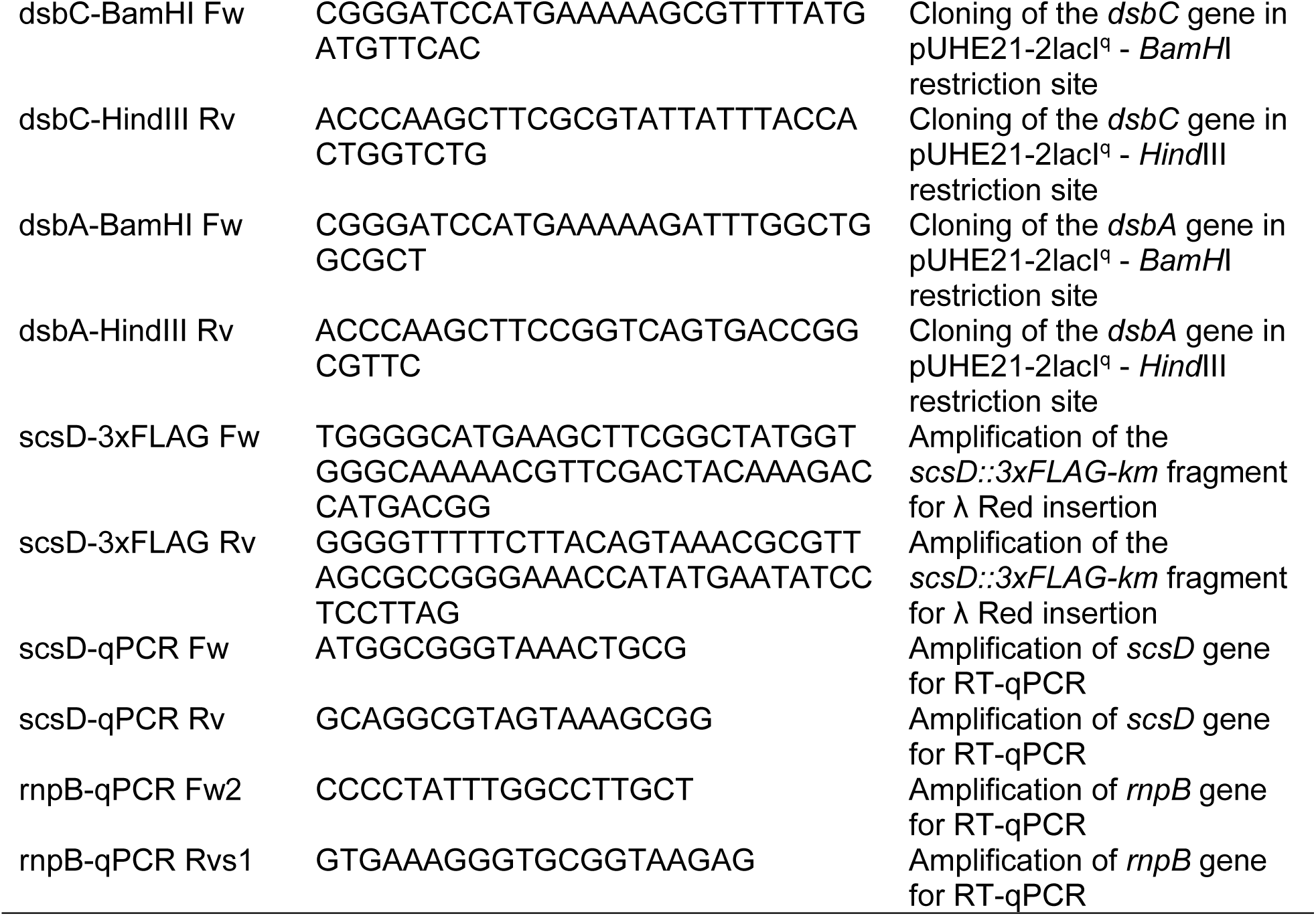
Oligonucleotides.

